# VoroCNN: Deep convolutional neural network built on 3D Voronoi tessellation of protein structures

**DOI:** 10.1101/2020.04.27.063586

**Authors:** Ilia Igashov, Kliment Olechnovic, Maria Kadukova, Česlovas Venclovas, Sergei Grudinin

## Abstract

**Motivation:** Effective use of evolutionary information has recently led to tremendous progress in computational prediction of three-dimensional (3D) structures of proteins and their complexes. Despite the progress, the accuracy of predicted structures tends to vary considerably from case to case. Since the utility of computational models depends on their accuracy, reliable estimates of deviation between predicted and native structures are of utmost importance.

**Results:** For the first time we present a deep convolutional neural network (CNN) constructed on a Voronoi tessellation of 3D molecular structures. Despite the irregular data domain, our data representation allows to efficiently introduce both convolution and pooling operations of the network. We trained our model, called VoroCNN, to predict local qualities of 3D protein folds. The prediction results are competitive to the state of the art and superior to the previous 3D CNN architectures built for the same task. We also discuss practical applications of VoroCNN, for example, in the recognition of protein binding interfaces.

**Availability:** The model, data, and evaluation tests are available at https://team.inria.fr/nano-d/software/vorocnn/.

## 1. Introduction

Protein structure prediction and protein structure analysis are very important problems in structural biology and bioinformatics. They have recently been subject to revolution thanks to multiple developments in several fields, most notably deep learning (1–3). Indeed, as the recent Critical Assessment of protein Structure Prediction (CASP) community-wide challenge has demonstrated, nowadays we are able to accurately predict protein structures even if they possess novel folds (4–8).

In the protein structure prediction field, so far deep-learning techniques have been routinely applied to regular two-dimensional data represented with matrices of multiple sequence alignments (2, 9, 10), or regular three-dimensional (3D) data of voxelized electron density maps (11–13). Given the unprecedented success of the former approaches in the general structure prediction task, it was a bit surprising to see that the latter could not achieve the same accuracy as more classical methods in the last CASP13 protein model quality assessment (MQA) exercise (14, 15). We believe that the data representation used in MQA methods that are based on 3D convolutional neural networks (CNN) is too complex for the currently available amount of data and computational resources. This work proposes a novel approach, called VoroCNN, that combines the advantages of versatility of 3D CNNs with a simpler data representation based on *Voronoi tessellation* of 3D space (16).

Proteins fold into specific three-dimensional (3D) structures as a result of interatomic interactions. Protein atoms interact among themselves and with the solvent, and these interactions rapidly decay with the distance. A rigorous way to define interatomic interactions is to construct a *Voronoi tessellation* of the protein atoms and relate every *Voronoi cell* to an atom and every *Voronoi cell face* to an interatomic contact (17–19). However, if a pair of contacting atoms is located near the surface of a protein structure, the corresponding Voronoi face may extend far away from the atoms. This problem can be circumvented by constraining the Voronoi cells of the atoms inside the boundaries defined by the solvent-accessible surface, enabling calculation of the areas for every atom-atom and atom-solvent contact. Such a solution has been implemented in Voronota (20), a software package specifically optimized to construct rapid tessellations for molecular structures, when the radii of balls (atoms) are not very different from each other. Each Voronoi tessellation-derived contact area describes the magnitude of the corresponding interaction. The relatively larger contact area indicates that the interaction is less overshadowed by adjacent interactions and vice versa. This trait, specific to the tessellation-based analysis of protein structures, naturally introduces non pairwise-additive molecular interactions. Interatomic contact areas as proxies for multibody interactions proved to be effective in various tasks, such as measuring deviation of models from the reference structure (21, 22) or the estimation of model accuracy in the absence of native structure (23–26).

Generally, it is rather difficult to operate on nonregular data structures in 3D space (27, 28). Therefore, to construct an efficiently *trainable* neural network, we decided to convert the initial 3D tessellations into *protein interaction graphs*. This allowed us to reuse all the knowledge already available for graph convolutional neural networks (29–33). We believe that a 3D tessellation can be reduced to a graph without loosing too much information. Indeed, relative coordinates in 3D space can be reconstructed from a set of pairwise distance observations, if we have a sufficient number of these, which are encoded as graph edges. This has been routinely demonstrated by various NMR-based techniques (34), and also recently by solving protein structures from residue contact maps (3, 35). Derivation of a protein interaction graph from Voronoi tessellation-based atomic contacts is then straightforward. Naturally, each protein atom corresponds to a *graph node*, each atom-atom contact - to a *graph edge*, and each contact area - to the weight of the edge. Every node can also contain additional tessellation-derived features. These can be the corresponding atom solvent-accessible area, the volume of the constrained Voronoi cell, etc. Also, such a graph representation inherently overcomes many orientation-dependence problems characteristic to some other methods.

## 2. Methods

The workflow of the VoroCNN method consists of the following steps. Firstly, given an atomistic 3D-model of a protein, we create the corresponding graph using the Voronoi 3D-tessellation method Voronota (20). Then, we convert the output of Voronota to a graph. After, we assign to the graph nodes some geometrical and physic-chemical features. Finally, we pass this graph as an input to a graph CNN that predicts local folding qualities of the input protein model. As we are building our graph using 3D geometric information about atom contacts, it was rather natural to us to choose the ground-truth local quality measure for the graph CNN that also assesses atomic contacts. Also, it has been recently shown that local measures, and contact area difference (CAD)-score in particular (21), are much more informative in multiple respects, and are also more consistent in selecting good models than global measures (36). Therefore, we have naturally chosen CAD-scores of each residue in the protein as the ground truth for the graph CNN. We should specifically mention that we primarily train the network to predict node-based scores, which are called local quality measures in the protein structure prediction community. Figure 1 shows a schematic representation of our method.

**Fig. 1.**
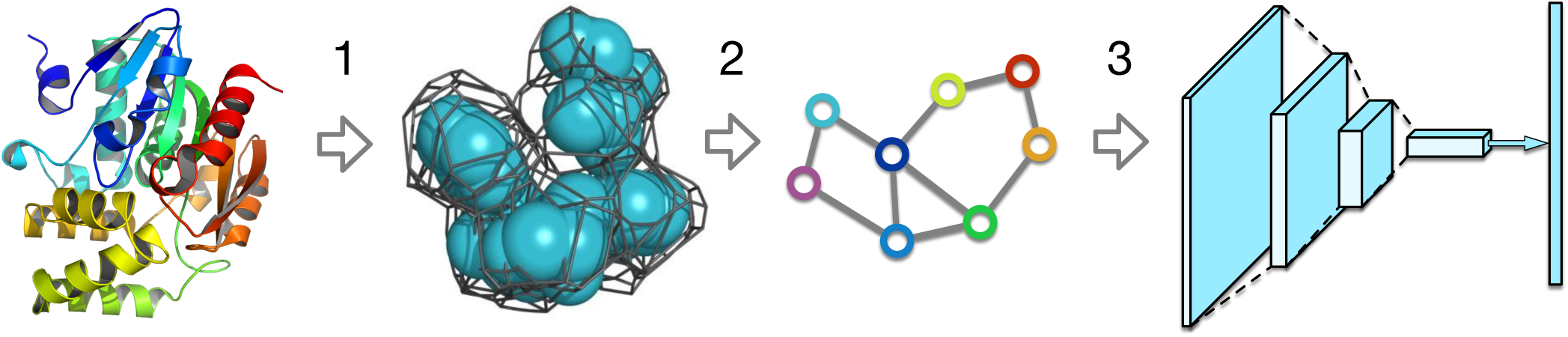
Schematic representation of the VoroCNN quality assessment method. Firstly, a Voronoi tessellation of a 3D-model is computed with Voronota. Then, based on Voronoi 3D-tessellation, a graph is built. Finally, a graph neural network predicts local CAD-scores of all residues in the initial model.

### A. Graph Representation

We represent the initial 3Dmodel of a protein as a *weighted unordered multigraph* with two types of edges, which are described in more detail below. The key property of the graph is that it implicitly keeps information about spatial relationship between the atoms based on the Voronoi 3D-tessellation of the protein model.

Nodes in the graph correspond to atoms in the protein structure. Each node of the graph contains a vector of features that describe the corresponding atom. These features include an atom type encoded with the one-hot representation (a binary vector with all “0”s and a single “1” value at the position corresponding to the type of the atom; we use 167 types in total following (11)), the solvent-accessible surface area for each atom computed with Voronota (20), the volume of atom’s Voronoi cell, also computed with Voronota, and the “buriedness” of the atom, which is a topological distance in the graph to the nearest solvent-accessible atom. We represent the whole set of nodes as a feature matrix **X** ∈ ℝ^*N*×*d*^, where *N* is the number of atoms in the protein structure and *d* = 170 is the dimension of the feature vector.

As we have mentioned above, our graph has two types of edges. Edges of the first type, the *contact edges*, correspond to a spatial relationship between the atoms. To construct these edges, we built a Voronoi partitioning on a set of balls whose positions and sizes are defined by the locations of the protein atoms and their van der Waals radii, correspondingly. We say that two atoms are *in a contact* if their Voronoi cells have non-zero contact surface. We consider that two atoms have a contact edge if these atoms are in a contact and there is no covalent bond between them. Edges of the second type, the *covalent edges*, correspond to the covalent bonds between the atoms that are in a contact. We should note that in some bad-quality models that contain atomic clashes, two atoms with a covalent bond may not be in a contact. In these cases we do not assign any edge to these atoms. We represent the two sets of edges as two *adjacency matrices*. For the contact edges we introduce a matrix **A**^c^ ∈ ℝ^*N*×*N*^, where *weights* 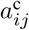 are equal to the area of the contact surface between Voronoi cells of the *i*-th and the *j*-th atoms if there is a contact edge between them, and zero otherwise. For the covalent edges we introduce a matrix **A**^b^ ∈ ℝ^*N*×*N*^, where *weights* 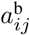 are equal to the area of the contact surface between Voronoi cells of the *i*-th and the *j*-th atoms if there is a covalent edge between them, and zero otherwise.

In order to improve numerical stability of the stochastic optimization and to add a certain level of regularization (30), we normalize the adjacency matrices according to the following equation,

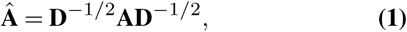

where **A** is an adjacency matrix and **D** is a diagonal matrix with nodes’ degrees at the diagonal. We only consider edges of the same type when computing degrees of the nodes. We also normalize the geometric features of the nodes, since the weights of the edges in our graph have a *geometric* origin, i.e. they are the contact surface areas. More precisely, the normalized features are the solvent-accessible surface areas and Voronoi cells’ volumes.

Finally, after the normalization, we split the covalent edges into 3 subtypes according to the types of covalent bonds, which can be single, double or aromatic. We also split the contact edges into 6 subtypes according to atoms’ sequence-separation distances. For example, the first type of edges is composed of atoms belonging to the same residue, the second type of edges is composed of atoms that are in two consecutive residues, etc. As a result, we obtain two adjacency tensors, 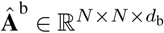 for the covalent edges with *d*_b_ = 3, and 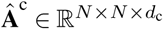 for the contact edges with *d*_c_ = 6.

### B. Graph Convolutional Neural Network

Here we introduce a graph neural network that solves the following problem. Given a graph of a protein model in atom-level representation, the aim is to predict the local CAD-scores (21) of all the residues in the model. The key components of our network are the convolutional and the pooling layers, which are described below.

#### B. Convolutional Layer

In the past years convolutional neural networks became very popular as an efficient method for various image processing tasks (37–40). The core idea that lies behind the convolutional layer is that it can learn local patterns that appear in different segments of the image. Very recently, similar approaches started to be adapted for graph structures (27, 33, 41). Contrary to images, graphs represent an irregular data domain that makes the definition of convolution operation on graphs more complicated. One common way to define the convolution operation on a graph is based on the idea that one should combine information about a graph node with information about its neighbors. This basic observation has been further developed into various realizations of convolutional layers for graphs (29–32). Today graph convolutional networks are becoming a popular alternative to more classical approaches in structural bioinformatics as well (42–46). This section introduces a graph convolutional layer that inherits from the same principles of sharing information between graph nodes and also takes into account the specificity of our data.

Our convolutional layer contains trainable tensors 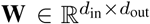, 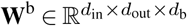, and 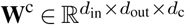, where *d*_in_ is the number of node features before passing them to the layer and *d*_out_ is the number of node features after the layer has been applied. Each layer transforms the feature matrix 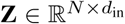 into 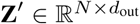 according to the following equation,

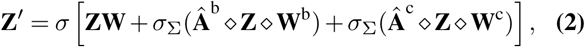

where the result **X** ◆ **Y** of the o operator is defined as

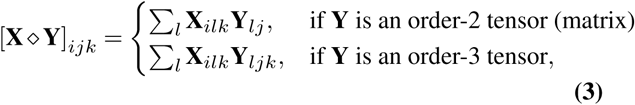

the function *σ*_Σ_ is defined as

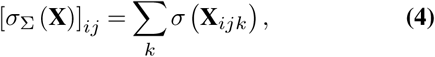

and *σ* is a nonlinear activation function. In the present work we use exponential linear unit (ELU) as the activation function (47).

#### B.2 Pooling Layer

Downsampling operations are often used in classical CNN architectures to reduce the data representation, achieve a better translational invariance, and extract hierarchical features. For images represented with pixels, downsampling can be implemented using convolutional filters with a stride that reduces the size of the output with respect to the input, or also using additional pooling layers that return one pixel according to an operation applied to several input pixels. However, downsampling in a graph is an open research problem, because it is unclear how to define this operation on a non-regular domain in the general case. Nonetheless, there are several approaches that are based on clustering algorithms (48–50). In this work we introduce a pooling operation that uses prior information about the topology and structure of the input graph.

Indeed, our graphs are very specific in sense that we have a strict hierarchy of the representation. More precisely, atoms in the input data are grouped into residues. This allows us to introduce a pooling layer that downsamples graph to the residue-level representation by averaging atoms’ feature vectors within each residue. After applying this layer, covalent edges become primitive, i.e. they simply represent the peptide chain of the protein. Therefore, after the pooling layer we keep working only with the contact edges. We should also specify that in this case the adjacency matrix is rewritten with the contact areas between the residues, which are computed as sums of the relevant inter-atom contact areas.

#### B.3. Network Architecture

We have tested and assessed multiple graph network architectures that are described in more detail below. The final architecture, VoroCNN, and its modification, VoroCNN-conv, are composed of a sequence of seven consecutive convolutional layers and one pooling layer in the middle of the sequence, as it is shown in Fig. 2. The first two convolutional layers are applied to the one-hot vectors only and reduce their length from 167 to 64. The next two convolutional layers are applied to the resulting nodes’ vectors concatenated with 3 atoms’ features. The pooling layer in the middle of the sequence downsamples the graph to the residue-level representation and the next three convolutional layers reduce the dimensionality of node features to 1. VoroCNN-conv has an additional 1D-convolutional layer at the very end.

**Fig. 2.**
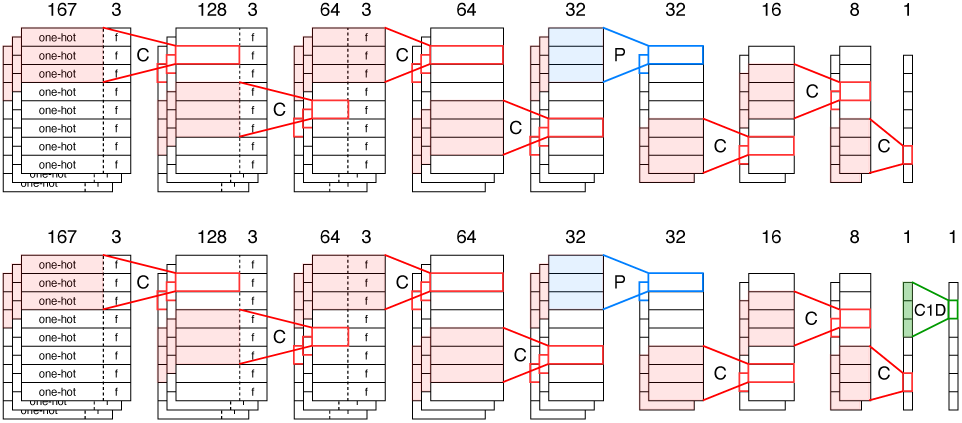
Architectures for the VoroCNN (top) and VoroCNN-conv (bottom) networks. Red color denotes the convolution operation (“C”), blue color denotes the pooling operation (“P”), green color denotes the 1D-convolution operation (“C1D”). Rectangles represent feature matrices, one row of a rectangle represents a feature vector of one graph node: it consists of one-hot part (marked as “one-hot”) and other features (marked as “f”). If there are no symbols inside the rectangles, it means that the convolution operation was applied to the whole vector and it is meaningless to distinguish one-hot and other features anymore. Multiple-stacked rectangles mean that the network operations are applied to the three-dimensional adjacency tensors. Numbers on top of the matrices correspond to the size of the feature vectors.

To train the network on local CAD-scores, we use Mean Squared Error (MSE) as a loss function,

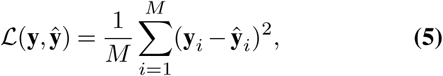

where **y** is a vector of ground-truth local CAD-scores, **ŷ** is a vector of VoroCNN predictions and *M* is a number of residues in a protein.

#### B.4 Training Parameters and Technical Details

To train the model we used the Adam optimizer with the learning rate of 0.001 (51). We stored and processed the adjacency matrices in the sparse format, and the whole training process was conducted in 15 parallel CPU threads. Thanks to the sparse representation of the data, the efficiency of the CPU training turned out to be slightly higher than that on the GPUs. Each thread in one iteration processed 300 models. The dataloader was organized in a way that the threads were sequentially fed with structural models corresponding to a particular target. Here we assume that each native structure, called a *target*, has multiple corresponding models. Only after processing of all the models for the current target the dataloader moved to another target. Such data-loading policy helped us to reduce the variance of the resulting models trained independently. One training iteration took on average 10 minutes on 15 Intel® Xeon® processors E5-2650 v2 @ 2.60GHz, and the models converge in about 10 iterations. The code was written in Python using the PyTorch library (52). All trained models, code, and preprocessing binaries are available at https://team.inria.fr/nano-d/software/vorocnn/.

### C. Datasets

To train and test our networks we used submissions from the previous CASP (Critical Assessment of protein Structure Prediction) challenges (4, 53). More precisely, for the training we used models from stage-2 server submissions of CASP[8-11] when tested on CASP12 submissions, and models from stage-2 server submissions of CASP[8-12] when tested on CASP13 submissions. We would like to mention that some of the CASP targets can form obligatory protein complexes, others can belong to membrane proteins, and also there can be targets with only low-quality models. To keep the physics of interactions, we tried to prune the training set as much as possible. Thus, we excluded from the training set some models and certain targets, as we explain below. Following our previous experiments (54), we have also augmented the set of input protein models by generating near-native conformations using the NOLB tool (55). Overall, our training dataset CASP[8-11] consisted of 333 target structures and 73418 models, and dataset CASP[8-12] consisted of 365 target structures and 79467 models. To test the performance of our network, we have also constructed two test datasets. The first included stage-2 server submissions of the CASP12 experiment (38 targets), the second included stage-2 server submissions of the CASP13 experiment (79 targets). We did not specifically filter out any models in this case. However, for both the training and the test sets, we also excluded models with unrecognized atom names, and models that contained hetero atoms. The full list of targets for the training and the test sets can be found in Supplementary Information (Tables S1 and S2).

#### C.1. Pruning the training set

In order to identify and exclude potential obligatory complexes and membrane proteins, and also to filter out targets that have only low-quality models, we performed the following procedure. For each of the targets, we computed its VoroMQA score (25) and retrieved the highest CAD-score (21) among all the models submitted for this target. Afterwards, we identified the targets corresponding to the low outliers of the VoroMQA scores and the low outliers of the maximum CAD-scores. The identified suspicious targets were then inspected visually and removed if deemed precarious. We also excluded from the training set those models that did not have all the residues of the corresponding target structures. Please see Table S1 from Supplementary Information for more details.

After the model filtering, we proceeded as follows. Firstly, we removed from the model all the excessive residues, i.e. those that were not present in the corresponding target structure. We also removed hydrogen atoms. Then, we computed per-residue ground-truth scores (local CAD-scores), and finally built the graphs.

#### C.2. Data Augmentation

The specificity of our training data is that it contains very few models with sufficiently-high CAD-scores. Indeed, more than 90% of the models in the training set have CAD-scores lower than 0.7. We have previously demonstrated that augmentation of the training data with good-quality models significantly improves the success rate of the predictions (54). Therefore, we reused this approach and generated random perturbation of the target structures using the nonlinear Normal Mode Analysis method NOLB (55). More precisely, we combined deformations along 100 slowest normal modes with random amplitudes. Then, we generated 50 decoy models for each target structure below the RMSD threshold of 0.9 Å. This threshold roughly corresponds to 0.95 global CAD-score.

## 3. Results and Discussion

### A. Test sets and metrics

We evaluated the performance of VoroCNN on CASP12 and CASP13 stage-2 datasets^1^. None of them were used for the training of the corresponding models. Our main goal was to assess the ability of VoroCNN to select the best model from a pool submitted for a certain target structure. This can be fulfilled in several ways, and in this work we report z-scores, ranks, per-target Pearson and Spearman correlations of modle scores (56) averaged over all the targets using the Fisher transformation (57). We provide results computed on the following metrics, the global CAD-score (21), the global lDDT-score (58), and GDT-TS (59, 60). We put emphasis on z-scores since they weight the predictions according to targets’ “difficulty”. In addition, z-score is the main assessment metric in the CASP challenges.

For the comparison with the state of the art, we have chosen several recent *single-model* methods. We want to specifically draw the reader’s attention to the fact that we only compare our results with the best single-model methods. The second class of methods, the *consensus-based* approaches (15), selects the best models based on the analysis of the whole pool of structures. They often perform better than the single-model methods, but can not assign a score to a single model without having access to the rest of the pool. Thus, we eliminated this class of methods from the comparison.

For the comparison we have chosen the Voronoi diagram-based method VoroMQA (25), the machine learning approach that uses orientational statistics of protein’s backbones SBROD (54), a descriptor-based method ProQ3 (61), which also has access to sequence information, and a 3DCNN approach Ornate (11). Since VoroCNN is trained to predict local per-residue scores, to obtain the global score of a model and compare with the other methods, we averaged the local predictions. We should also add that we specifically designed a variation of VoroCNN, called VoroCNN-conv, with an additional smoothing layer trained to smooth the local scores along the sequence.

### 2. VoroCNN results

Here we report results of VoroCNN and VoroCNN-conv obtained on CASP[12-13] stage-2 datasets. The numbers on CASP12 for SBROD were computed by us for a previous publication (54). We also locally computed the corresponding numbers for Ornate. For ProQ3 and VoroMQA, we used results from the server submissions archive downloaded from the official CASP website. For CASP13, we obtained the results of SBROD, VoroMQA-B and ProQ3, ProQ3D-lDDT from the official CASP website as well. We should mention that SBROD did not estimate several targets in CASP13, so we assigned “-2” as SBROD’s z-scores for these targets, to be on par with the official CASP assessment policy. When calculating SBROD’s mean correlations and ranks, we excluded these targets from the analysis. We also used ground-truth CAD-scores, lDDT-scores and GDT-TS scores from the CASP website.

Table 1 lists the results for CASP12. Here, the VoroCNN models were trained on CASP[8-11] data. Table 2 compares results for CASP13. Here, the VoroCNN models were trained on CASP[8-12] data. For each of these two cases we independently launched 8 training processes and then chose the *best* trained model achieved on the test data with respect to the CAD z-scores. We can see that in both benchmarks VoroCNN and VoroCNN-conv outperform other methods if the aim is to predict global CAD-scores. For the CASP12 benchmark, we outperform the other methods in CAD, lDDT and GDT-TS z-scores and also in CAD and lDDT ranks. For the CASP13 benchmark, we outperform the other methods CAD z-scores and CAD ranks. However, our method is not the best according to the correlation measure, especially in the GDT-TS case. This behavior can be explained by the fact that although we average local predictions in order to get the global score, in reality global CAD-scores (and even more GDT-TS) are not a simple average of the local predictions. On the other hand, it is obvious that for high-quality models all the local scores should be high and should also correlate with the global score. Therefore, ability of a local-scoring algorithm to select high-quality models correctly does not imply that the same algorithm also correctly predicts global scores in general. Since local scores show different properties from those of global scores (such as GDT-TS), optimizing the performance according to one type of scores unavoidably makes the results worse according to the other type (36).

**Table 1.** Comparison of VoroCNN, VoroCNN-conv, Ornate, VoroMQA, SBROD, and ProQ3 on the CASP12 stage-2 dataset.

| Method | z-score | | | rank | | | Pearson's $r$ | | | Spearman's $\rho$ | | |
| --- | --- | --- | --- | --- | --- | --- | --- | --- | --- | --- | --- | --- |
|  | CAD | IDDT | GDT-TS | CAD | IDDT | GDT-TS | CAD | IDDT | GDT-TS | CAD | IDDT | GDT-TS |
| ProQ3 | 1.670 | 1.441 | 1.141 | 11.961 | 13.500 | 25.961 | 0.801 | **0.775** | 0.692 | 0.750 | **0.734** | **0.615** |
| SBROD | 1.282 | 1.234 | 1.034 | 23.579 | 18.842 | 27.329 | 0.762 | 0.726 | **0.694** | 0.685 | 0.670 | 0.581 |
| VoroMQA | 1.410 | 1.178 | 0.761 | 17.171 | 18.158 | 36.421 | 0.803 | 0.759 | 0.638 | 0.766 | 0.725 | 0.561 |
| Ornate | 1.780 | 1.440 | 1.180 | 10.776 | 13.355 | 24.539 | **0.828** | 0.729 | 0.573 | 0.781 | 0.686 | 0.499 |
| VoroCNN | **1.871** | **1.518** | **1.191** | **9.276** | **12.000** | 25.829 | 0.817 | 0.704 | 0.565 | 0.774 | 0.682 | 0.509 |
| VoroCNN-conv | **1.857** | **1.480** | **1.271** | **8.197** | **11.447** | **19.408** | 0.823 | 0.700 | 0.563 | **0.783** | 0.679 | 0.508 |

**Table 2.** Comparison of VoroCNN, VoroCNN-conv, Ornate, VoroMQA-B, SBROD, ProQ3-lDDT, and ProQ3 on the CASP13 stage-2 dataset.

| Method | z-score | | | rank | | | Pearson's $r$ | | | Spearman's $\rho$ | | |
| --- | --- | --- | --- | --- | --- | --- | --- | --- | --- | --- | --- | --- |
|  | CAD | IDDT | GDT-TS | CAD | IDDT | GDT-TS | CAD | IDDT | GDT-TS | CAD | IDDT | GDT-TS |
| ProQ3 | 1.457 | 1.210 | 0.980 | 18.494 | 22.804 | 33.715 | 0.771 | 0.731 | 0.640 | 0.732 | 0.717 | 0.595 |
| ProQ3-IDDT | 1.495 | **1.257** | **1.023** | 15.620 | **20.873** | 30.044 | **0.832** | **0.782** | **0.712** | 0.792 | **0.773** | 0.666 |
| SBROD | 1.052 | 0.894 | 0.734 | 18.321 | 21.850 | **28.979** | 0.772 | 0.724 | 0.673 | 0.762 | 0.753 | **0.670** |
| VoroMQA-B | 1.363 | 1.185 | 0.976 | 21.513 | 23.696 | 33.816 | 0.802 | 0.776 | 0.661 | 0.762 | 0.731 | 0.608 |
| Ornate | 1.410 | 1.134 | 0.843 | 19.051 | 26.525 | 40.127 | 0.814 | 0.752 | 0.606 | 0.781 | 0.714 | 0.577 |
| VoroCNN | **1.581** | 1.149 | 0.948 | **13.625** | 27.042 | 36.333 | 0.763 | 0.671 | 0.541 | 0.728 | 0.630 | 0.531 |
| VoroCNN-conv | **1.508** | 1.219 | 1.004 | **15.297** | 21.810 | 29.778 | **0.832** | 0.756 | 0.648 | **0.798** | 0.734 | 0.610 |

### C. Local scores

As our model predicts *per-residue* folding qualities, these can be visually illustrated in all major molecular visualization systems. Figure 3A-B provides a visual comparison of VoroCNN predictions for the crystallographic structures, their CASP13 models, and also ground-truth local CAD-scores. Figure 3A shows a structure and models of strigolactone receptor (pdb code 5CBK), which consists of multiple alpha-helices. Figure 3B demonstrates a beta-propeller structure and models of Wdr5 protein (pdb code 2O9K). One can see that the predictions of VoroCNN for the native structures and the corresponding models are visually very similar to the ground truth.

**Fig. 3.**
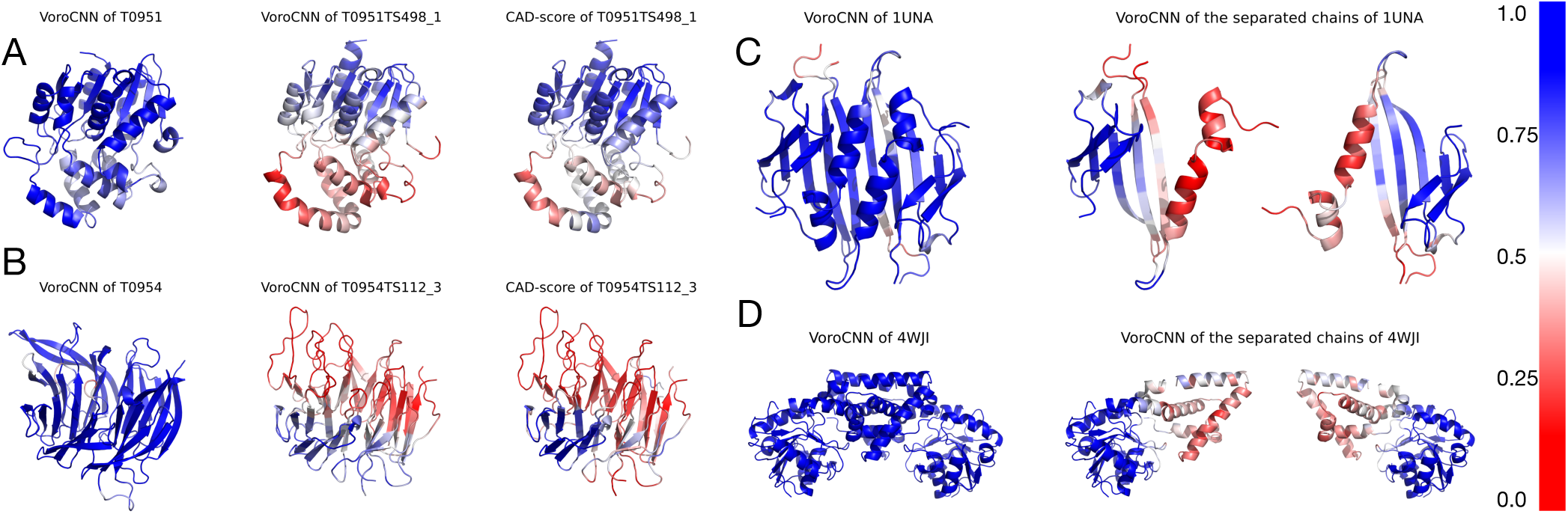
Illustration of local scores predictions. The color-bar on the right corresponds to the values of the scores. A) VoroCNN predictions for the T0951 CASP target (left), T0951TS498_1 model (center), and the ground-truth local CAD-scores (right). B) VoroCNN predictions for the T0954 CASP target (left), T0954TS112_3 model (center), and the ground-truth local CAD-scores (right). C) VoroCNN predictions for the obligatory complex of bacteriophage RNA-binding protein (pdb code 1UNA, left) and its individual subunits (right). D) VoroCNN predictions for the obligatory complex of cyclohexadienyl dehydrogenase (pdb code 4WJI, left) and its individual subunits (right).

A generally interesting question is how much predictions of the local scores can be useful for the structural bioinformatics community. An obvious application of the local scores, as it is demonstrated in Fig. 3A-B, is to highlight local structural inaccuracies in protein models. Indeed, it has been recently demonstrated that local scores, and CAD-score in particular, are well suited for promoting physical realism of protein models (36). Local scores are also more robust in dealing with multi-domain structures as well as movements of loops or local structural motifs. Another practical example can be an analysis of protein binding interfaces. Figure 3C-D shows VoroCNN predictions for two obligatory complexes and their individual subunits. We can clearly see that the binding interfaces have lower scores compared to the rest of the structure, and are very visually distinguishable. This can be explained by the specificity of these interfaces. Very often they are hydrophobic and it is energetically unfavorable for them to be exposed into solvent. This is why they can be detected by the local-scoring schemes, e.g., VoroCNN.

### D. Binary classification

An interesting general question is how well VoroCNN can distinguish target (native) structures from the models. Let us consider here VoroCNN as a binary classifier, meaning an algorithm that predicts one of two classes that a given model belongs to. In order to evaluate the quality of VoroCNN binary classification, we used global scores predicted by VoroCNN for all models and targets from our test sets CASP12 and CASP13^2^. Figure 4 (left) shows distributions of the global scores predicted for targets and models from CASP12 by VoroCNN trained on CASP[8-11]. Figure 4 (right) shows the same distributions for CASP13 by VoroCNN trained on CASP[8-12]. In both cases we can see a clear separation between the two distributions. For CASP12 scores, the value of ROC-AUC, the area under the ROC-curve (62), equals to 0.942. For CASP13 scores, ROC-AUC is 0.953. Thus we can conclude that VoroCNN has learned to discriminate target structures from the models.

**Fig. 4.**
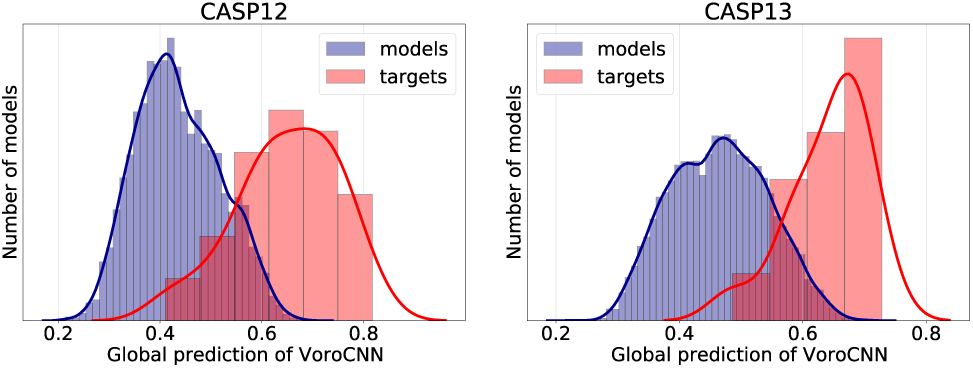
Distribution of VoroCNN scores on target structures and models from CASP12 (left) and CASP13 (right). Solid lines represent kernel density estimations of the corresponding distributions.

### E. Tested architectures

In order to design the final architecture we studied a number of variations of the network. This section briefly describes them and discusses their benefits and drawbacks, a schematic representation of the networks can be found in Fig. S1 from Supplementary Information. Figure 5 summarizes averaged trajectories of correlation coefficients and z-scores of CAD-scores on the test data for the selected network designs. For models trained on CASP[8-11] the metrics are measured on CASP12, and for models trained on CASP[8-12] the metrics are measured on CASP13. Evaluation on other metrics, box-plot results, and comparison with the state-of-the-art methods can be found in Figs. S2-S7 from Supplementary Information.

**Fig. 5.**
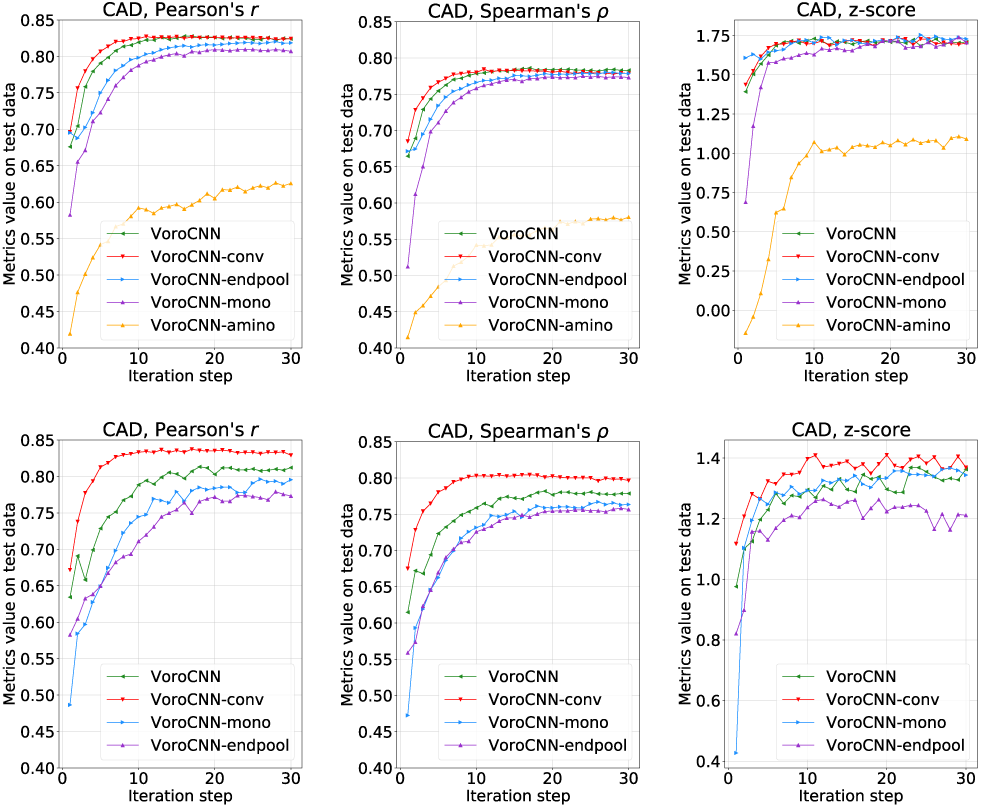
Mean trajectories of CAD Pearson and Spearman correlations and z-scores of VoroCNN model and its modifications. The first row represents averaged results of models trained on CASP[8-11] and tested on CASP12. The second row shows averaged results of models trained on CASP[8-12] and tested on CASP13.

At the very beginning we started with a more classical architecture named VoroCNN-endpool that differs from the final VoroCNN in the position of the pooling layer. In the VoroCNN-endpool architecture, the pooling is applied at the very end of the network and plays solely a technical role – it aggregates atom-scores into residue-scores. The average performance of VoroCNN-endpool network on CAD zscores is similar to the performance of VoroCNN. However, VoroCNN-endpool has lower correlation scores, which can be explained by the fact that in VoroCNN-endpool residue-level scores are not optimized after pooling. Moreover, VoroCNN-endpool has more parameters compared to the final architecture, since the pooling is applied at the very end, and, as a result, it also requires more time to train.

Another interesting effect that we noticed is connected to the structure of the input data. As we have described above, we represent contact- and covalent-edge adjacency matrices as three-dimensional tensors. Precisely, we expand twodimensional matrices along the third dimension by the subtypes of the edges. According to eq. Eq. (2), it means that we have separate trainable matrices for each edge subtype. We also tested an approach when the graph edges are represented by a single matrix. This simplified architecture, called VoroCNN-mono, performed worse on average. However, the number of its trainable parameters is dramatically reduced compared to VoroCNN.

In order to improve results of our method on the GDTTS metrics, we tried to smooth the local predictions of VoroCNN. To do that, we added a trainable 1D-convolutional layer at the very end of the network. Indeed, the designed VoroCNN-conv architecture performs better than VoroCNN on GDT-TS. Moreover, except for the CAD-scores, VoroCNN-conv beats VoroCNN in all metrics on CASP13 (see Table 2).

Finally, we have also tested the VoroCNN-amino architecture that processes only residue-level data. Surprisingly enough, it demonstrated somewhat less impressive results compared to the recently published residue-level methods (45, 46). This can be partially explained by its design. We have only tested a very simple network that contains 4 sequential convolutional layers.

Along with the described designs, we have also tried various shapes and numbers of convolutional layers, played with the loss function, learning rates and feature normalization approaches. For example, we have tested weighted MSE with weights corresponding to the ground-truth z-scores or ground-truth global CAD-scores as alternative loss functions. We have also varied the network depth. If we increase the depth of the base architecture by adding more convolutional layers, there is no significant improvement in the network performance. However, reducing of the network depth results in worse CAD z-scores. Thus, VoroCNN and VoroCNN-conv are the final designs of all the tested networks.

## 4. Conclusion

This work suggests a novel way to learn on 3D protein folds and 3D macromolecular data in general. For the first time we demonstrate the applicability of learning on 3D Voronoi tessellations using graph convolutional networks. Our results confirm a high potential of using 3D tessellation and graph representation in general in various learning tasks in structural bioinformatics. Indeed, despite the complexity of the VoroCNN model and a rather big number of free parameters, our results are comparable to the state of the art and better than those of the very recent 3D CNN architectures trained on regular volumetric representations, e.g. Ornate. Thus, we believe that currently, given the available amounts of training data and computational resources, Voronoi tessellation is a better representation of 3D protein structure than raw volumetric data.

This work also illustrates a potential of methods that predict local folding accuracies for various structural bioinformatics applications. Indeed, we have demonstrated that VoroCNN can highlight structural inaccuracies in protein models, and can also distinguish protein binding interfaces.

## Acknowledgements

The authors would like to thank Elodie Laine from Sorbonne Université for the discussions during the study and proofreading the manuscript.

## Supplementary Information

**Fig. S1.**
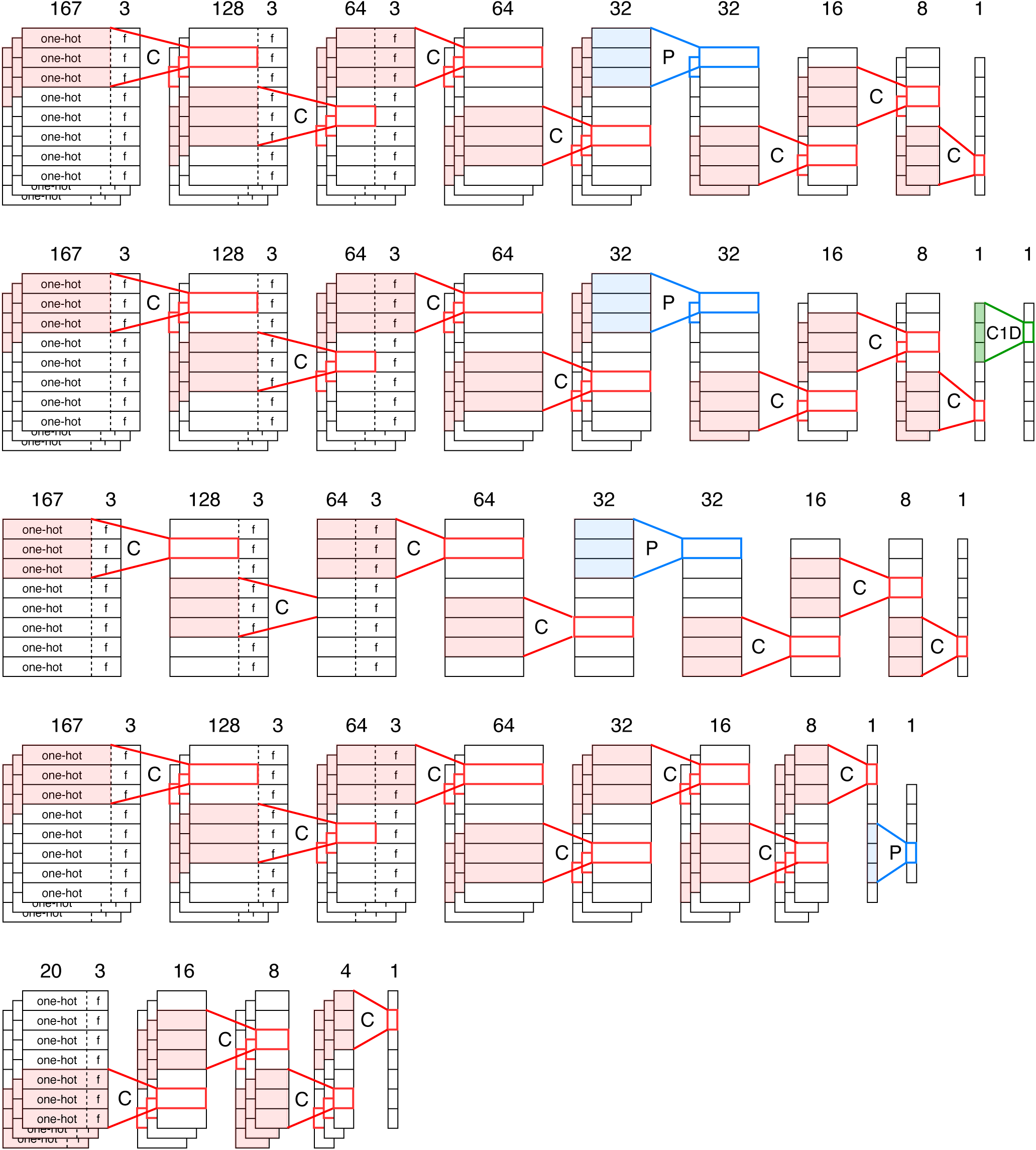
Schematic representation of tested network architectures (from top to bottom): VoroCNN, VoroCNN-conv, VoroCNN-mono, VoroCNN-endpool and VoroCNN-amino. Red color denotes the convolution operation (≪C≫), blue color denotes the pooling operation (≪P≫), and green color denotes the 1D-convolution operation (≪C1D≫). Rectangles represent feature matrices, one row of a rectangle represents a feature vector of one graph node: it consists of one-hot part (marked as ≪one-hot≫) and other features (marked as ≪f≫). If there are no symbols inside a rectangle, it means that the convolution operation was applied to the whole vector and it is meaningless to distinguish one-hot and other features anymore. Multiple stacked rectangles mean that the network operations are applied to three-dimensional adjacency tensors. Numbers on top of the matrices correspond to the size of the feature vectors.

**Fig. S2.**
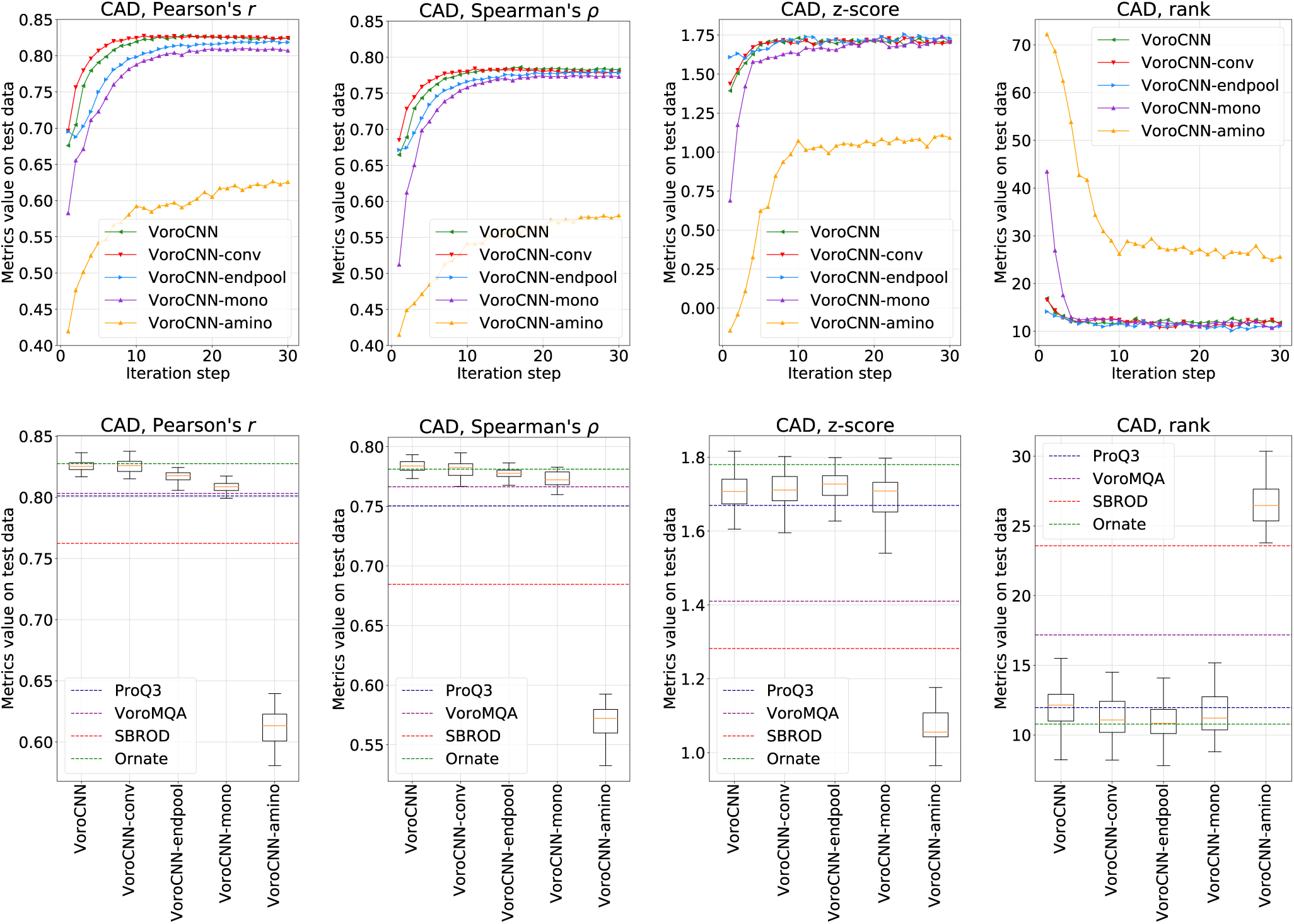
Trajectories (top) and boxplots (bottom) of Pearson and Spearman correlations, z-scores and ranks on CAD-scores evaluated on CASP12 for models trained on CASP[8-11]. Trajectories are averaged over launches (for each architecture we independently trained and evaluated 8 models). For the boxplots we used all the results of each step of the training of each model beginning from the 20th iteration, assuming that from this iteration step the quality of predictions did not change from iteration to iteration significantly.

**Fig. S3.**
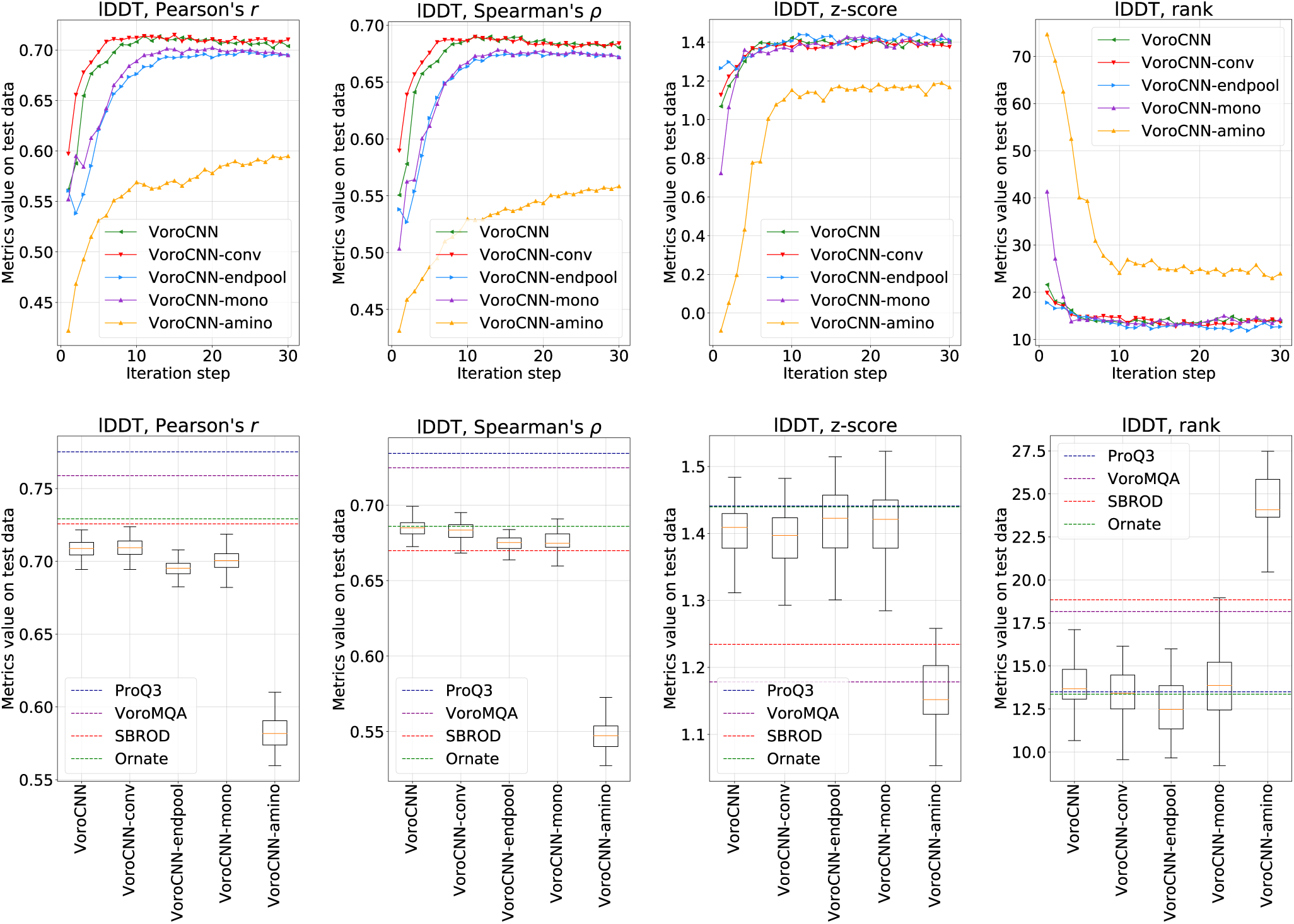
Trajectories (top) and boxplots (bottom) of Pearson and Spearman correlations, z-scores and ranks on lDDT-scores evaluated on CASP12 for models trained on CASP[8-11]. Trajectories are averaged over launches (for each architecture we independently trained and evaluated 8 models). For the boxplots we used all the results of each step of training of each model beginning from the 20th iteration, assuming that from this iteration step the quality of predictions did not change from iteration to iteration significantly.

**Fig. S4.**
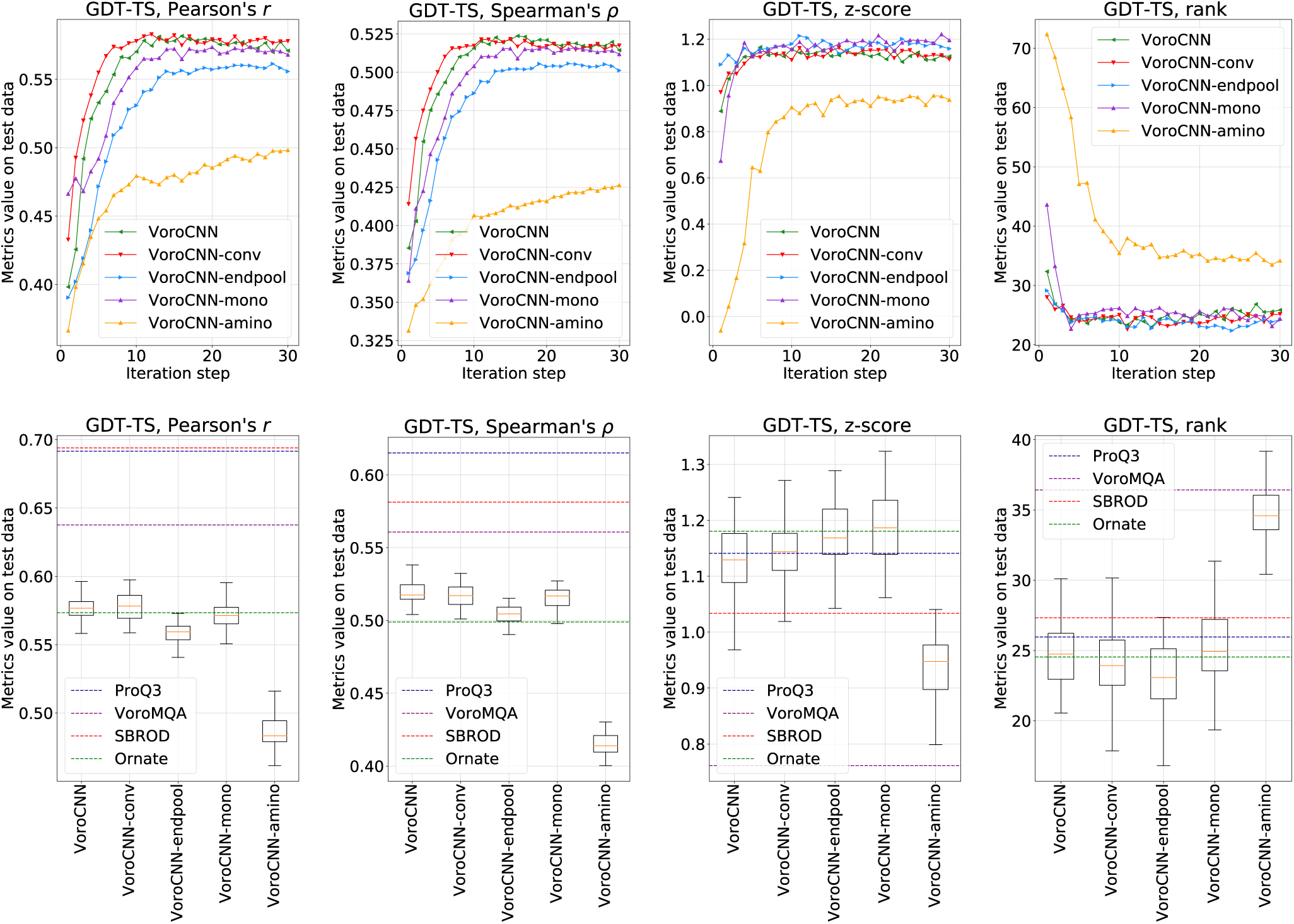
Trajectories (top) and boxplots (bottom) of Pearson and Spearman correlations, z-scores and ranks on GDT-TS evaluated on CASP12 for models trained on CASP[8-11]. Trajectories are averaged over launches (for each architecture we independently trained and evaluated 8 models). For the boxplots we used all the results of each step of the training of each model beginning from the 20th iteration, assuming that from this iteration step the quality of predictions did not change from iteration to iteration significantly.

**Fig. S5.**
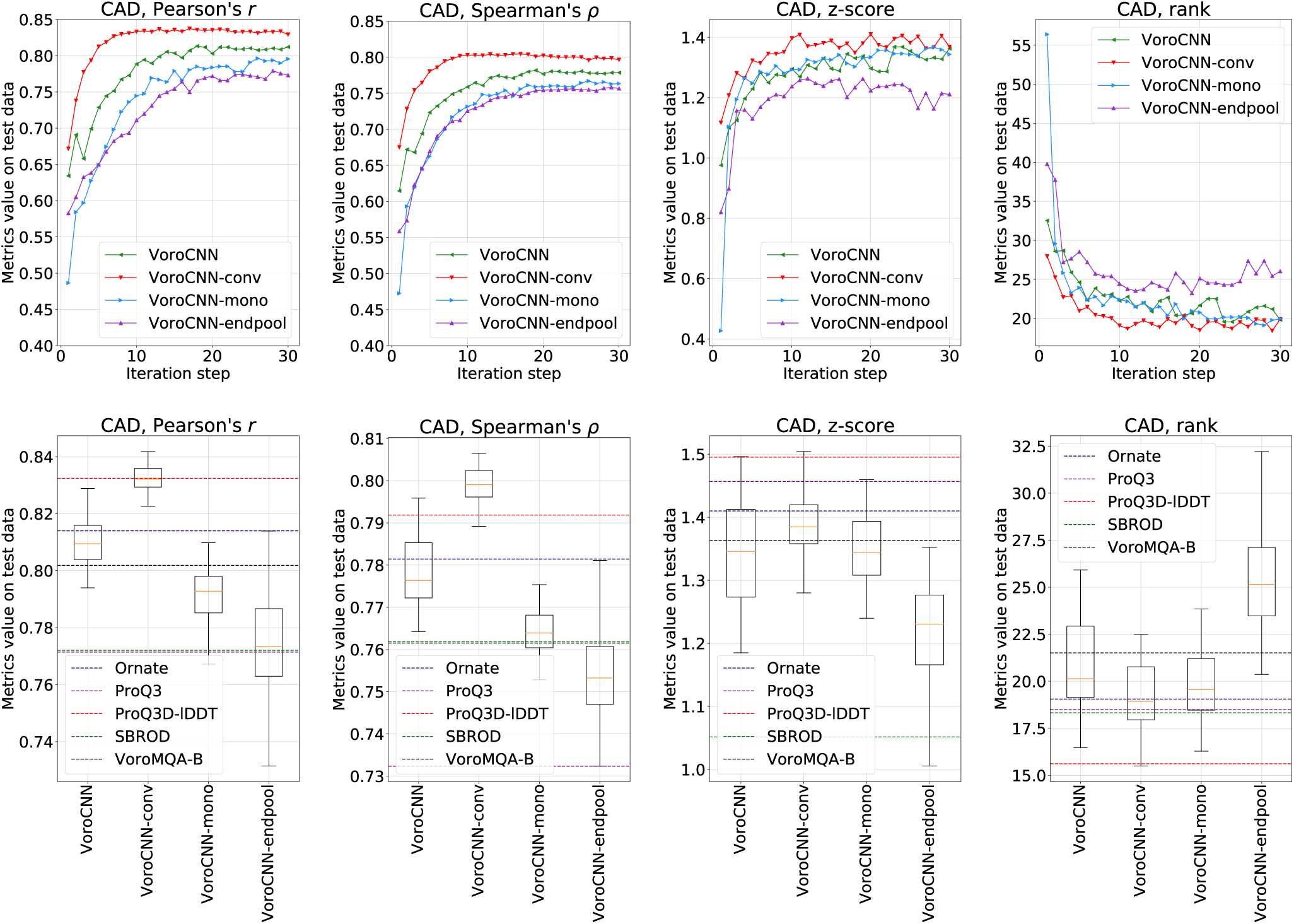
Trajectories (top) and boxplots (bottom) of Pearson and Spearman correlations, z-scores and ranks on CAD-scores evaluated on CASP13 for models trained on CASP[8-12]. Trajectories are averaged over launches (for each architecture we independently trained and evaluated 8 models). For the boxplots we used all the results of each step of the training of each model beginning from the 20th iteration, assuming that from this iteration step the quality of predictions did not change from iteration to iteration significantly.

**Fig. S6.**
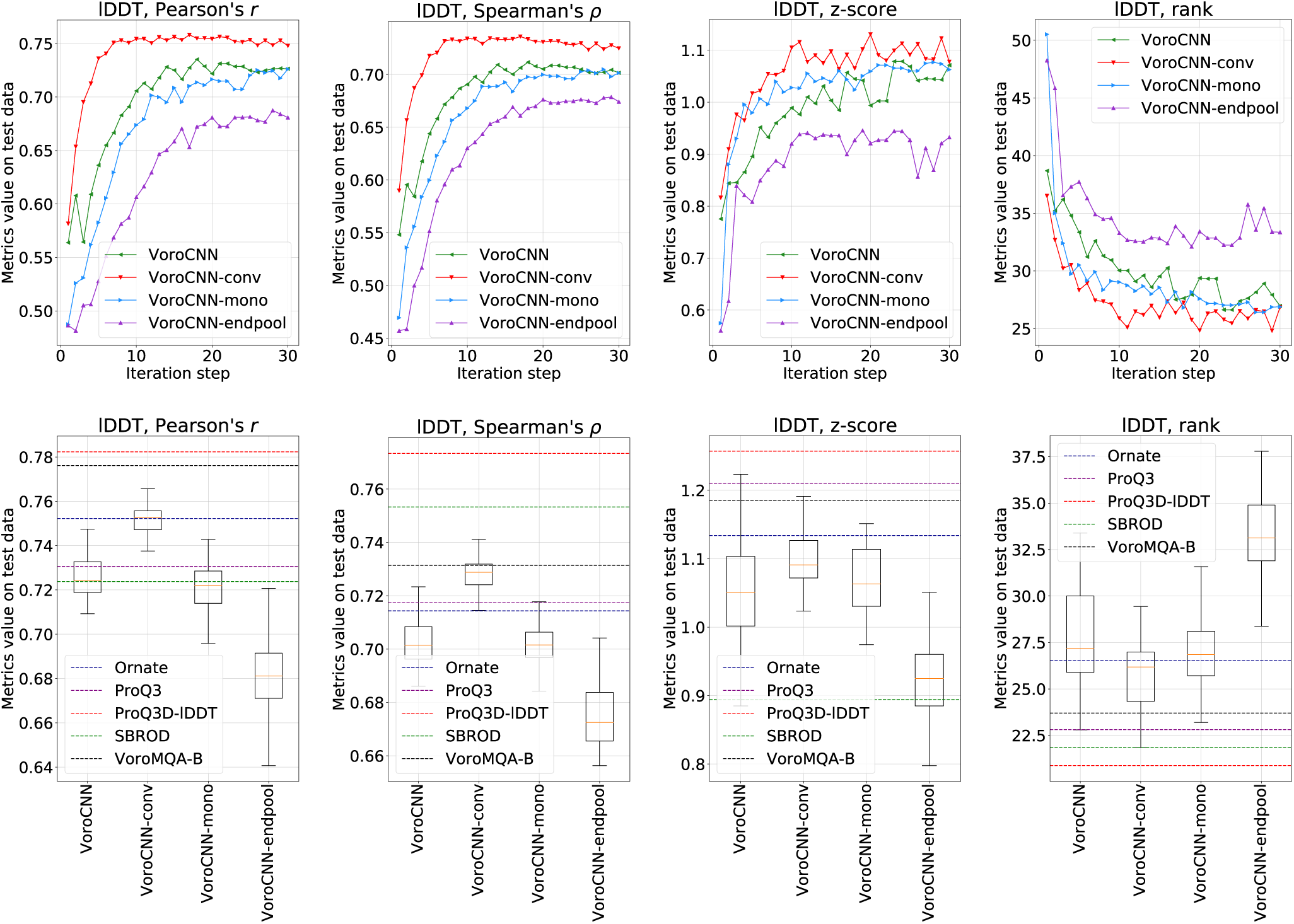
Trajectories (top) and boxplots (bottom) of Pearson and Spearman correlations, z-scores and ranks on lDDT-scores evaluated on CASP13 for models trained on CASP[8-12]. Trajectories are averaged over launches (for each architecture we independently trained and evaluated 8 models). For the boxplots we used all the results of each step of the training of each model beginning from the 20th iteration, assuming that from this iteration step the quality of predictions did not change from iteration to iteration significantly.

**Fig. S7.**
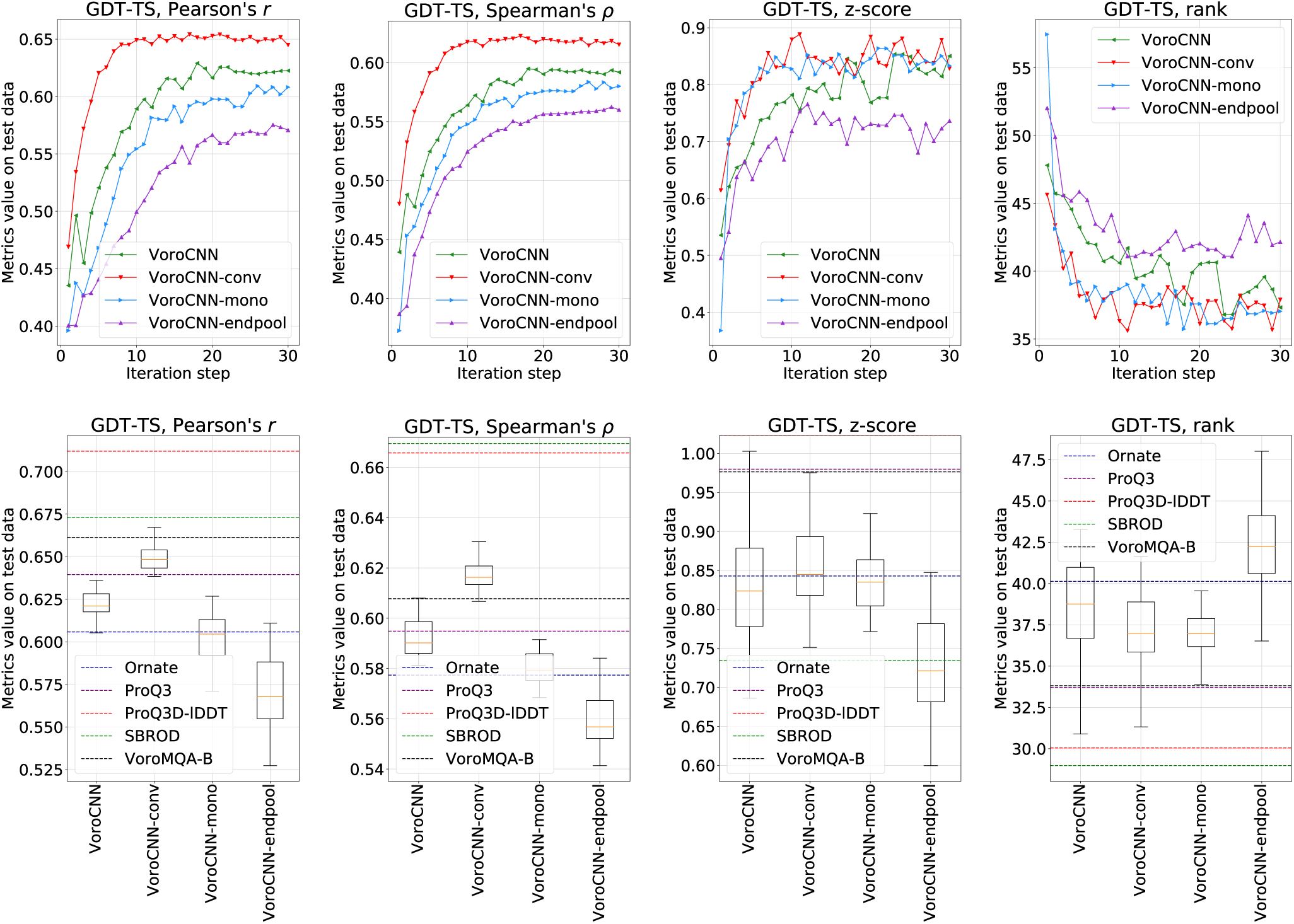
Trajectories (top) and boxplots (bottom) of Pearson and Spearman correlations, z-scores and ranks on GDT-TS evaluated on CASP13 for models trained on CASP[8-12]. Trajectories are averaged over launches (for each architecture we independently trained and evaluated 8 models). For the boxplots we used all the results of each step of the training of each model beginning from the 20th iteration, assuming that from this iteration step the quality of predictions did not change from iteration to iteration significantly.

**Table S1.**
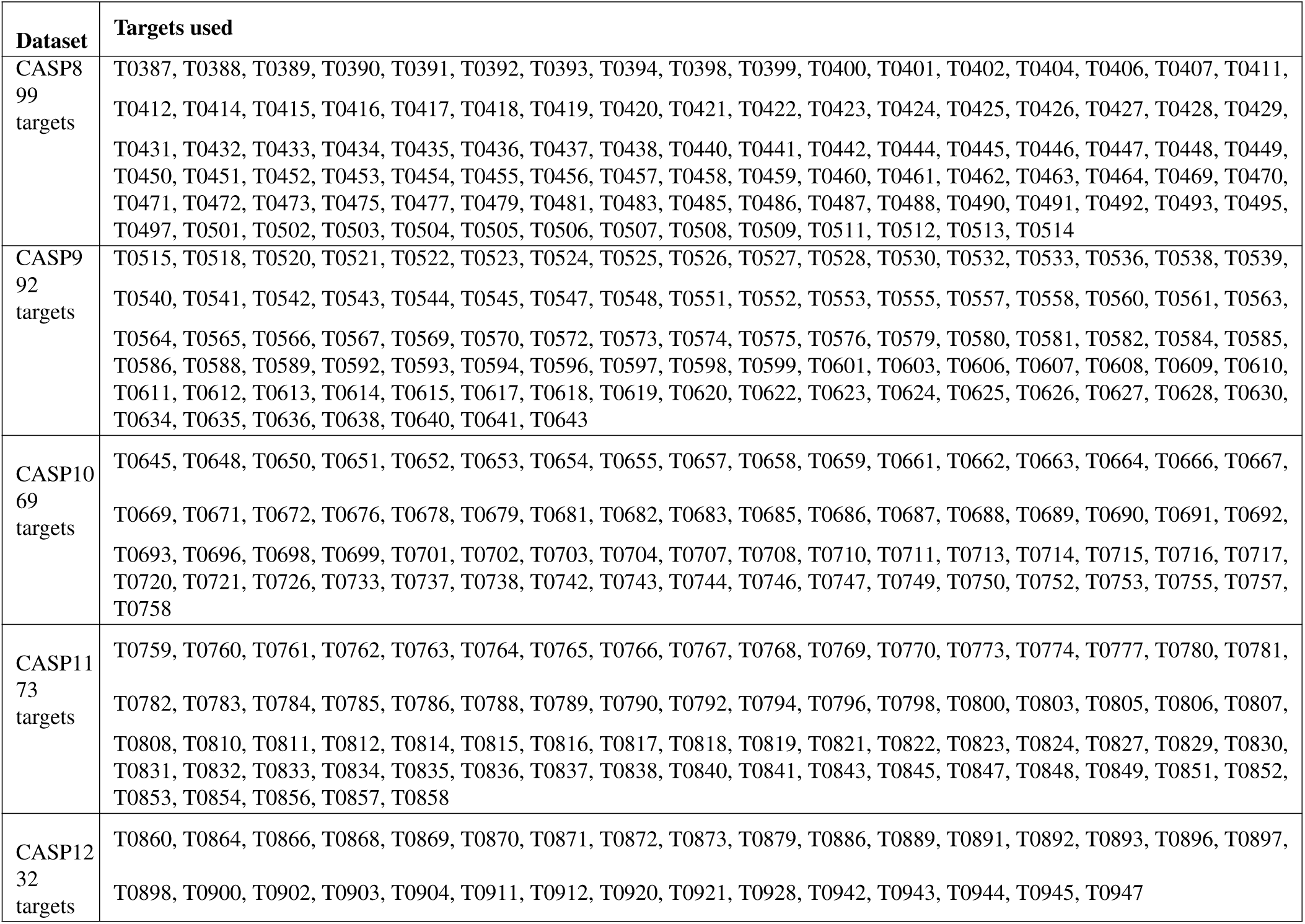
CASP targets that are present in the training datasets.

| Dataset | Targets used |
| --- | --- |
| CASP8<br>99<br>targets | T0387, T0388, T0389, T0390, T0391, T0392, T0393, T0394, T0398, T0399, T0400, T0401, T0402, T0404, T0406, T0407, T0411, T0412, T0414, T0415, T0416, T0417, T0418, T0419, T0420, T0421, T0422, T0423, T0424, T0425, T0426, T0427, T0428, T0429, T0431, T0432, T0433, T0434, T0435, T0436, T0437, T0438, T0440, T0441, T0442, T0444, T0445, T0446, T0447, T0448, T0449, T0450, T0451, T0452, T0453, T0454, T0455, T0456, T0457, T0458, T0459, T0460, T0461, T0462, T0463, T0464, T0469, T0470, T0471, T0472, T0473, T0475, T0477, T0479, T0481, T0483, T0485, T0486, T0487, T0488, T0490, T0491, T0492, T0493, T0495, T0497, T0501, T0502, T0503, T0504, T0505, T0506, T0507, T0508, T0509, T0511, T0512, T0513, T0514 |
| CASP9<br>92<br>targets | T0515, T0518, T0520, T0521, T0522, T0523, T0524, T0525, T0526, T0527, T0528, T0530, T0532, T0533, T0536, T0538, T0539, T0540, T0541, T0542, T0543, T0544, T0545, T0547, T0548, T0551, T0552, T0553, T0555, T0557, T0558, T0560, T0561, T0563, T0564, T0565, T0566, T0567, T0569, T0570, T0572, T0573, T0574, T0575, T0576, T0579, T0580, T0581, T0582, T0584, T0585, T0586, T0588, T0589, T0592, T0593, T0594, T0596, T0597, T0598, T0599, T0601, T0603, T0606, T0607, T0608, T0609, T0610, T0611, T0612, T0613, T0614, T0615, T0617, T0618, T0619, T0620, T0622, T0623, T0624, T0625, T0626, T0627, T0628, T0630, T0634, T0635, T0636, T0638, T0640, T0641, T0643 |
| CASP10<br>69<br>targets | T0645, T0648, T0650, T0651, T0652, T0653, T0654, T0655, T0657, T0658, T0659, T0661, T0662, T0663, T0664, T0666, T0667, T0669, T0671, T0672, T0676, T0678, T0679, T0681, T0682, T0683, T0685, T0686, T0687, T0688, T0689, T0690, T0691, T0692, T0693, T0696, T0698, T0699, T0701, T0702, T0703, T0704, T0707, T0708, T0710, T0711, T0713, T0714, T0715, T0716, T0717, T0720, T0721, T0726, T0733, T0737, T0738, T0742, T0743, T0744, T0746, T0747, T0749, T0750, T0752, T0753, T0755, T0757, T0758 |
| CASP11<br>73<br>targets | T0759, T0760, T0761, T0762, T0763, T0764, T0765, T0766, T0767, T0768, T0769, T0770, T0773, T0774, T0777, T0780, T0781, T0782, T0783, T0784, T0785, T0786, T0788, T0789, T0790, T0792, T0794, T0796, T0798, T0800, T0803, T0805, T0806, T0807, T0808, T0810, T0811, T0812, T0814, T0815, T0816, T0817, T0818, T0819, T0821, T0822, T0823, T0824, T0827, T0829, T0830, T0831, T0832, T0833, T0834, T0835, T0836, T0837, T0838, T0840, T0841, T0843, T0845, T0847, T0848, T0849, T0851, T0852, T0853, T0854, T0856, T0857, T0858 |
| CASP12<br>32<br>targets | T0860, T0864, T0866, T0868, T0869, T0870, T0871, T0872, T0873, T0879, T0886, T0889, T0891, T0892, T0893, T0896, T0897, T0898, T0900, T0902, T0903, T0904, T0911, T0912, T0920, T0921, T0928, T0942, T0943, T0944, T0945, T0947 |

**Table S2.** CASP targets that are present in the test datasets.

| Dataset | Targets used |
| --- | --- |
| CASP12<br>38<br>targets | T0859, T0860, T0862, T0863, T0864, T0866, T0868, T0869, T0870, T0871, T0872, T0873, T0879, T0886, T0889, T0891, T0892, T0893, T0896, T0897, T0898, T0900, T0902, T0903, T0904, T0911, T0912, T0918, T0920, T0921, T0922, T0928, T0941, T0942, T0943, T0944, T0945, T0947 |
| CASP13<br>79<br>targets | T0949, T0950, T0951, T0953s1, T0953s2, T0954, T0955, T0957s1, T0957s2, T0958, T0959, T0960, T0961, T0962, T0963, T0964, T0965, T0966, T0967, T0968s1, T0968s2, T0969, T0970, T0971, T0973, T0974s1, T0974s2, T0975, T0976, T0977, T0978, T0979, T0980s1, T0980s2, T0981, T0982, T0983, T0984, T0985, T0986s1, T0986s2, T0987, T0989, T0990, T0991, T0992, T0993s1, T0993s2, T0995, T0996, T0997, T0998, T1000, T1001, T1002, T1003, T1004, T1005, T1006, T1008, T1009, T1010, T1011, T1013, T1014, T1015s1, T1015s2, T1016, T1017s1, T1017s2, T1018, T1019s1, T1019s2, T1020, T1021s1, T1021s2, T1021s3, T1022s1, T1022s2 |

## Footnotes

1 We used only models submitted by servers.

2 Here, for CASP13, we used only 13 publicly available target structures.

